# Parental thermal conditions affect the brain activity response to alarm cue in larval zebrafish

**DOI:** 10.1101/2024.06.02.597016

**Authors:** Jade M. Sourisse, Julie L. Semmelhack, Celia Schunter

## Abstract

With temperature being a crucial factor affecting the physiology of ectothermic animals, global warming will likely impact neural mechanisms aquatic organisms use to perceive their environment over generations. However, exposure to elevated temperature during specific life stages and across generations may confer fish resilience through phenotypic plasticity. In this study, we investigate the effects of developmental and parental temperature on brain activity response to an olfactory cue in the larval zebrafish, *Danio rerio*. We exposed parents during reproduction and their offspring during development to control (28°C) or elevated temperature (30°C) and observed the response of the larval telencephalon to an alarm cue using live calcium imaging. Parental exposure to elevated temperature decreased the time till maximum brain activity response regardless of the offspring’s developmental temperature, revealing that parental thermal conditions can affect the excitability of the offspring’s neural circuitry. Furthermore, brain activity duration was affected by the interaction between parental and offspring thermal conditions, tending to last longer when either parents or offspring were exposed to elevated temperature, yet more similar to control when elevated temperature was experienced by both parents and offspring. This could represent an anticipatory parental effect influencing the offspring’s brain response to match the parental environment, or an early developmental effect occurring within a susceptible short time window post-fertilization. Overall, our results suggest that future predicted warming can alter processes involved in brain transmission and show that parental conditions could aid in the preparation of their offspring to respond to olfactory stimuli in a changing environment.

## Introduction

Temperature is arguably the principal environmental factor that can influence fish performance (Brett, 1971; Johnston and Dunn, 1987) and has known effects on a variety of behaviours (Fukuhara, 1990; Stoner, Ottmar and Hurst, 2006; Biro, Beckmann and Stamps, 2010) and brain function (Szabo *et al*., 2008; Andreassen *et al*., 2022). As global warming is experienced in aquatic environments at an unprecedented rate (Allen *et al*., 2018; Pörtner *et al*., 2019), we may expect near-future thermal conditions to alter crucial processes in fish, including sensory responses. Olfaction is one such crucial sensory modality in fish that is crucial for behaviours such as feeding, migration and predation escape (v. Frisch, 1938; Hara, 1975). It is mediated by olfactory sensory neurons (OSNs) in the olfactory epithelia, which converge in the telencephalon in a region called the olfactory bulb (Laberge and Hara, 2001). From the olfactory bulb, second order projections to other telencephalic olfactory areas provide input to neuronal pathways involved in behavioural responses such as escape or freezing (Jesuthasan and Mathuru, 2008; Miyasaka *et al*., 2009). The olfactory circuitry is sensitive to higher temperatures and shows temperature dependent hyperexcitability in fish (Flerova and Gdovskii, 1975; Døving and Belghaug, 1977) indicating potential implications for the persistence of olfactory triggered behavioural responses under global warming. However, it is unclear how near-future thermal conditions will impact olfactory processing in the fish brain (Tigert and Porteus, 2023). Therefore, understanding how warming modulates brain functioning is crucial to better predict the effects of rapid climate change on fish behaviour and survival.

One way by which fish may show resilience to global warming is phenotypic plasticity, which is the capacity of an individual to adjust its phenotype to changing conditions without altering its genetic constitution (Crozier and Hutchings, 2014; Fox *et al*., 2019). Throughout development, specific windows of susceptibility exist during which organisms are more responsive to external conditions (Burggren and Mueller, 2015), such as early embryonic stages in fishes (Mueller *et al*., 2015; Flynn and Todgham, 2018; Melendez and Mueller, 2021). For instance, temperature during fish embryogenesis is known to influence later metabolic performance and behaviour into adulthood (Jonsson and Jonsson, 2014, 2018). Furthermore, a plastic response to global warming may occur not only within, but across generations when parents alter the phenotype of their offspring via non-genetic inheritance, named intergenerational plasticity (O’Dea *et al*., 2016; Fox *et al*., 2019). In fish, parental exposure can provide increased acclimation to elevated temperature to their offspring (Donelson *et al*., 2012; Shama *et al*., 2014). Parental effects of warming were shown to affect the brain, including expression of genes expression involved in neuromuscular junction development (Bernal *et al*., 2022) and hormonal pathways (Veilleux, Donelson and Munday, 2018). Despite the importance of those acclimation mechanisms to temperature elevation, how such within and intergenerational effects of warming may affect the offspring’s brain function response is not well understood, due to the complexity of the vertebrate brains. Understanding the mechanisms behind the changes caused by global warming in fish responses to environmental cues is therefore challenging.

To investigate the potential parental contributions to thermal acclimation in the fish brain as it senses its environment, we can make use of the zebrafish model. The neuronal circuitry governing olfactory-mediated processes has been extensively described in this model species (Friedrich, Jacobson and Zhu, 2010) thanks to the engineering of genetically encoded calcium indicators that allow brain activity to be observed at the larval stage, when the brain has functional yet less complex neuronal networks than the adult stage (Kettunen, 2012). Furthermore, temperature conditions experienced by parents were shown to confer metabolic compensation in their offspring as a transgenerational response in this species (Massey and Dalziel, 2023). Finally, as the olfactory fear response is innate in zebrafish (Jesuthasan and Mathuru, 2008) it can be observed in its simplest form without the need for learning experiences to occur. Antipredator behaviour in zebrafish can be olfactory-mediated: if they perceive an olfactory alarm cue coming from an injured conspecific in their surrounding environment, they reduce or even suppress their locomotion activity (Speedie and Gerlai, 2008). Therefore, we can observe an innate olfactory response of ecological relevance in the larval stage.

In our study, we investigate the effects of developmental elevated temperature on the brain activity response of zebrafish larvae to an alarm olfactory cue, the Conspecifics Alarm Cue (CAC). We hypothesize that future predicted thermal conditions for the end of the century will impact the neuronal circuitry by provoking hyperexcitability. Furthermore, we also assess the relative influence of parental thermal conditions and very early embryonic temperature exposure to elevated temperature on the offspring’s brain activity response to CAC. We expect that previous exposure to elevated temperature will alter brain activity patterns in response to CAC of offspring reared under near-future predicted conditions. By characterizing thermally induced brain activity changes in response to CAC, we aim at determining how future predicted temperature will affect the environmental perception of fish through the olfactory neuronal circuitry.

## Methods

### Animals, housing, and temperature exposure

Zebrafish breeders (5 females, 1 male; *Tg(elavl3:Has.H2B-GcaMP6s)* strain) were obtained from the Hong Kong University of Science and Technology zebrafish husbandry facilities. This zebrafish strain is unpigmented and possesses a genetically encoded calcium indicator, allowing to show the calcium ion levels increase in the brain during neuronal activity. The parent breeders were housed in recirculating systems (80 x 37 x 32 cm), with a 14/10h light-dark cycle. They were fed twice a day with TetraMin flakes. pH and nitrate levels were measured weekly using a Seven2GO portable pH meter (Mettler Toledo) and a HI97728 nitrate photometer (Hanna Instruments), respectively. Parent breeders were first reared in control conditions (28°C) for 46 days, during which fertilized eggs were collected to constitute an offspring group from “control’ parental conditions (later named P28). Then, parents were exposed to elevated temperature (30°C, reached over a day) and bred from the next seven days on to collect fertilized eggs belonging to an elevated temperature group (later named P30), for which warming was experienced during spawning and early embryogenesis. We selected the control temperature as it is the optimal rearing temperature of zebrafish in laboratory settings and chose the treatment temperature as +2°C as current IPCC scenarios estimate that because of climate change temperatures will globally increase between 1.5°C and 2°C above pre-industrial levels by the end of the century (Allen *et al*., 2018; Pörtner *et al*., 2019). Temperature was measured, adjusted, and recorded every 60 seconds with heaters (Schego) and a STC-1000 Thermostat (Elitech). Temperatures experienced by the parents differed significantly before and during the “heatwave” (Wilcoxon test, p-value = 1.743×10^-8^): the treatment temperature over the experimentation period was 29.8 ± 0.2°C, whereas the mean control temperature was 27.7 ± 0.2°C (Table S1; Fig. S1).

For both groups of offspring according to parental spawning temperature (P28 or P30), fertilized eggs (at the end of the cleavage period of development; Kimmel et al., 1995) were collected in the morning and then placed in Petri dishes filled with Danieau’s solution embryo medium (17 mM NaCl, 2 mM KCl, 0.12 mM MgSO_4_, 1.8 mM, Ca(NO_3_)_2_, 1.5 mM HEPES). They were housed in in DSI-060D incubators (Digisystem). Offspring from both elevated temperature groups were themselves divided into two thermal groups, either control (28°C, later named O28) or elevated (30°C, later named O30) temperature. Temperature inside the incubators was automatically adjusted every 60 seconds. The embryo medium was changed daily. Embryos were reared under a 14/10h light-dark cycle and from five days post fertilization (dpf) onwards they were fed a larval diet (Zeigler Bros) daily until seven dpf. Temperatures between offspring treatments differed significantly (Wilcoxon test, p-value = 3.883×10^-10^): the “treatment” temperature over the experimentation period was 28.1 ± 0.2°C, whereas the mean control temperature was 30.1 ± 0.1°C (Table S1; Fig. S2). This study was carried out in approval of the Committee on the Use of Live Animals in Teaching and Research (CULATR) of the University of Hong Kong (#5614-21 and #22-257).

### Brain imaging data acquisition

At 7 dpf, larvae were embedded in low-melting agarose inside a small Petri dish (Ø = 45mm, h = 16mm) filled with 6mL of medium and let to acclimate for two to three hours. After acclimation to embedding, larvae were placed under a confocal laser scanning microscope (LSM 980; Carl Zeiss) that recorded brain activity from the top using an excitation light of 488nm (FITC, Oregon Green) in the telencephalic region. Larvae were first exposed to 0.5mL of Danieau solution as a control introduced in the Petri dish by using a 2.5 mL syringe. After a 5min interval, they were then exposed to 0.5mL of Conspecifics Alarm Cue (CAC), a substance that is known to trigger an alarm response such as reduced or suppressed locomotion (Jesuthasan and Mathuru, 2008). Donor zebrafish larvae reared together with the subjects were sacrificed by head concussion to produce the CAC, at a concentration of 4 donors/mL and homogenizing their bodies in water with a sterile mortar and pestle, 5 min or less before cue exposure time (Sourisse *et al*., 2023). A total of 37 larvae were then used to record the brain activity response to CAC exposure (for the P28-O28 group, n = 8; for the P28-O30 group, n = 10; for the P30-O28 group, n = 9; for the P30-O30 group, n = 10). The confocal microscope recorded a total of 20 pictures per run, with each pictures shot every 1.65 seconds. Therefore, the acquisition of the brain activity response for each individual lasted 33 seconds.

### Statistical analyses

Stack time series acquired were opened in Fiji v 2.14.0 (Schindelin *et al*., 2012) and stacks were aligned using the SIFT plug in (Saalfeld and Tomancák, 2008). To measure fluorescence, the telencephalon was manually defined as a discrete Region of Interest (ROI) in each frame. Its plane area was directly measured from the aligned stack acquisition (μm^2^) as a proxy of brain development (Table S2). Differences in fluorescence (ΔF) were then calculated as follows: the baseline fluorescence value (F) of the ROI was chosen as the lowest value during the first 1-3 frames of the time series, which corresponds to a period of low activity. To calculate ΔF throughout the acquisition, F was subtracted to the corresponding ROI’s fluorescence value at any given time following the baseline frame (ΔF). Values of ΔF and ΔF/F over time were first used to compare runs performed on the same larva (control vs CAC) ensuring that the brain activity response recorded during exposure to CAC produced a curve of positive values of ΔF over time whereas in the case of the control, ΔF would not produce a positive peak but decrease in ΔF over time due to fluid addition (Fig. 1 as example). Then, ΔF/F values obtained for CAC in the four different thermal groups were used to assess the effect of parental or offspring temperature on the brain activity response to CAC. Specifically, several metrics were statistically compared in R (R Core Team, 2018) such as the maximum ΔF/F value in the run (maxΔF/F), the time at which maxΔF/F occurred (maxΔF/F time) and the brain activity response length (defined as duration for which ΔF> 0; Table S2). For maxΔF/F, one outlier was removed (Grubbs test, p-value = 0.019) before comparison. A two-way ANOVA was chosen when the response variable showed homogeneity of variances as per tested with a Bartlett’s test, otherwise a Scheirer-Ray-Hare test was performed.

**Figure 1:**
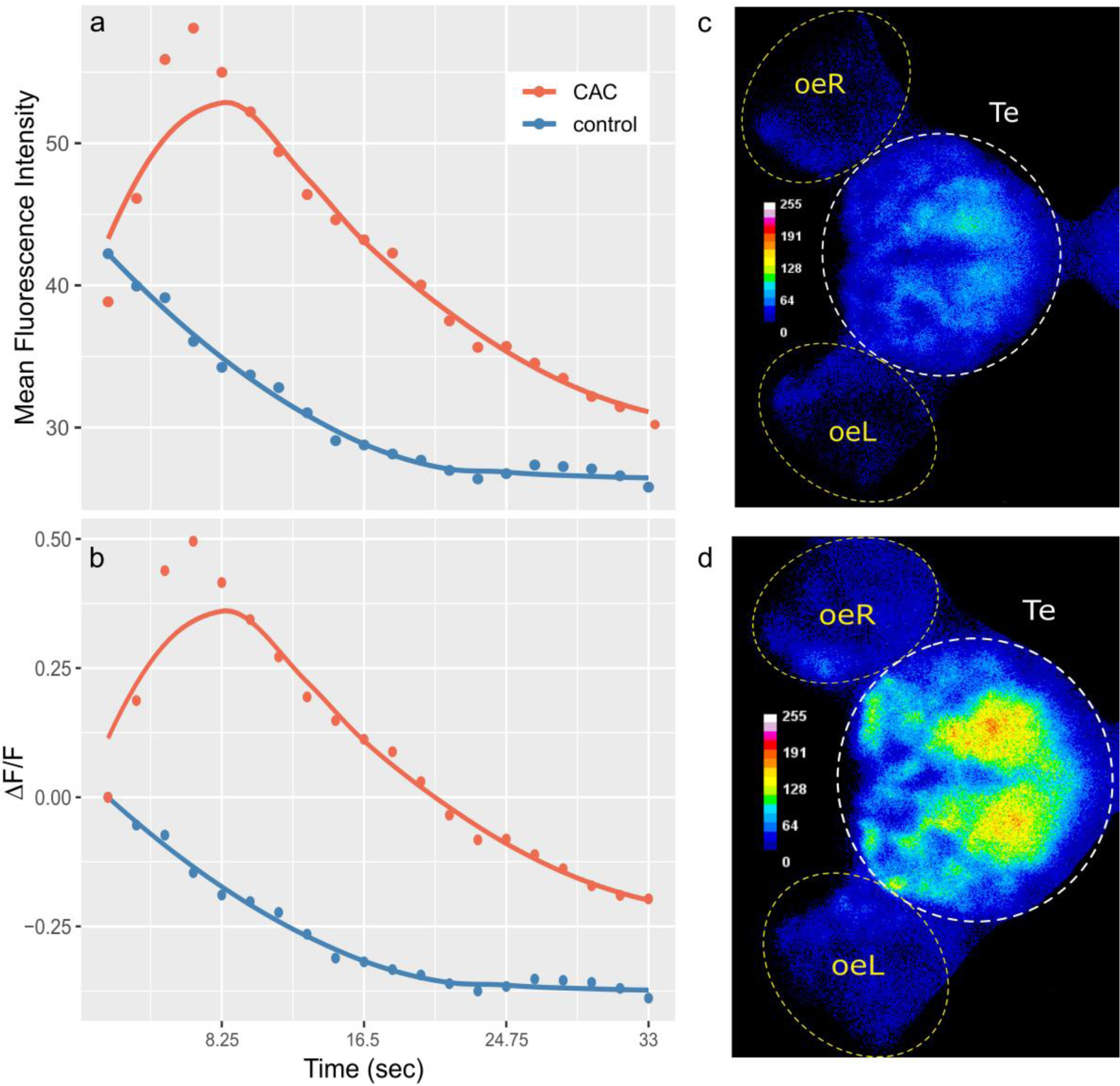
(a-b) Mean Fluorescence Intensity (a) and ΔF/F (b) over time of the telencephalon region of individual #N+30-2-89 as it is exposed to control Danieau solution (blue) or Conspecifics Alarm Cue (CAC, red); dots correspond to the measured values and lines plot smoothed conditional means; (c-d) images of forebrain activity during control Danieau solution (c, frame 1) or CAC (d, frame 4) exposure; the colour bars plot fluorescence intensity for each pixel; oe= olfactory epithelium; L = left; R = right; Te = Telencephalon

## Results

We hypothesized that warming would alter the brain activity response of zebrafish larvae responding to CAC. Surprisingly, there was no effect of parental nor offspring temperature on the maximum brain activity level maxΔF (Fig. S3), nor on maxΔF/F (Fig. 2) which accounts for individual variances in fluorescence baseline levels in the brain. This means that neuronal response in the forebrain did not increase with parental nor offspring exposure to elevated temperature during the olfactory response.

**Figure 2:**
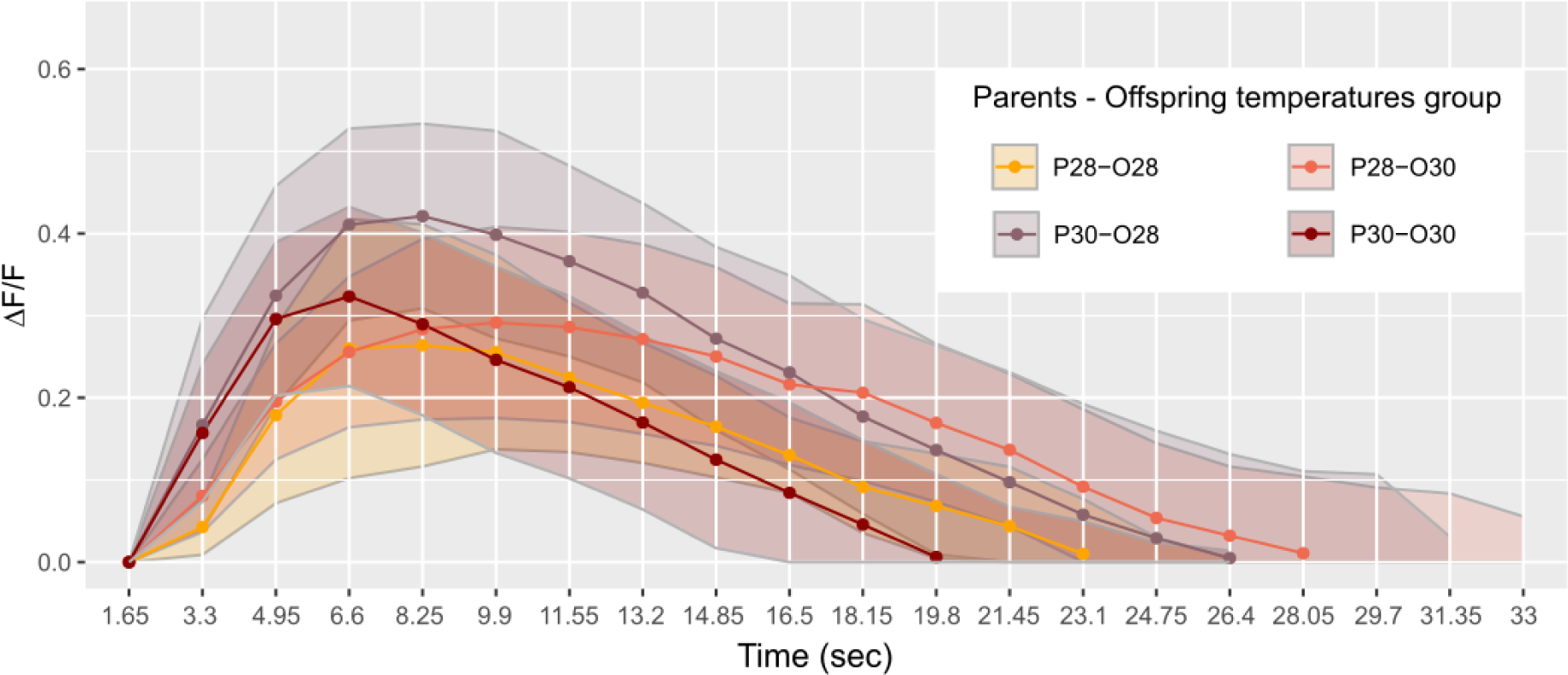
Mean telencephalon calcium fluorescence ratio (ΔF/F) over time (frame) in zebrafish larvae exposed to Conspecifics Alarm Cue (CAC); parents that produced these offspring experienced either control (P28) or elevated temperature (P30) conditions; larvae were reared in either control (O28) or elevated temperature (O30) conditions; dots represent mean values whereas ribbons around solid lines correspond to standard deviation

However, there was a significant effect of the parental spawning temperature on maxΔF/F time (Scheirer-Ray-Hare test, p-value = 0.0097; Fig. 3a). Larvae of parents who bred at control temperature reached their maximum brain activity at 9.53 ± 3.13 seconds (n = 18) whereas larvae born from parents experiencing elevated temperature reached maximum activity earlier, at 7.29 ± 1.68 seconds (n = 19).

**Figure 3:**
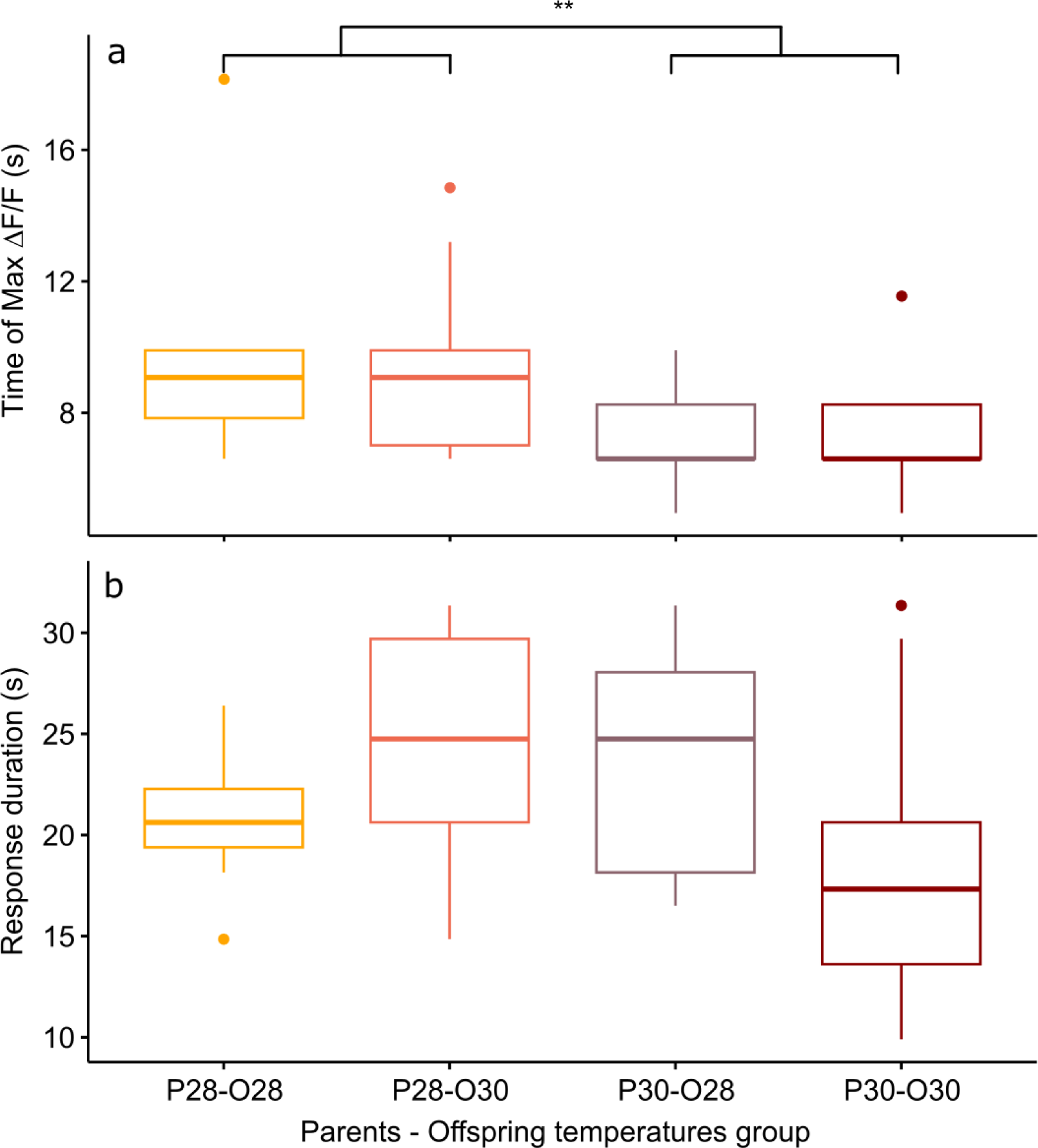
Mean time of maximum calcium fluorescence (time of max ΔF/F; a) and mean duration (b) of the telencephalon response to CAC in zebrafish larvae exposed to Conspecifics Alarm Cue (CAC); parents that produced these offspring experienced either control (P28) or elevated temperature (P30) conditions; larvae were reared in either control (O28) or elevated temperature (O30) conditions; stars (**) in (a) indicates the significant difference between offspring from the P28 group and that of the P30 group

Development in elevated temperature had an effect on the variability of maxΔF/F time (Bartlett test, p=value = 0.0123) with offspring reared at elevated temperature (n = 20) reaching their maximum brain activity level at different times compared to offspring reared at control temperature (n= 17) for which maximum brain activity levels were mostly reached around 8.44 ± 2.9 seconds. However, there was no effect of developmental temperature on the time of maxΔF/F itself (Scheirer Ray Hare test, p-value = 0.852) nor any interaction effect between parental spawning and offspring thermal conditions (Scheirer Ray Hare test, p-value = 0.934).

Although the brain response duration was not affected by parental spawning temperature (two-way ANOVA, p-value = 0.278), nor by offspring thermal exposure (two-way ANOVA, p-value = 0.686), there was an interaction effect of those two factors (two-way ANOVA, p-value = 0.022), suggesting that developmental warming has a different effect on zebrafish larvae brain activity depending on the thermal conditions experienced either during early embryogenesis and/or by their parents at time of reproduction (Fig. 3b). Larvae of parents exposed to control temperature and themselves also reared at control temperature had a brain activity response duration of 20.83 ± 3.63 seconds (n = 8). This response duration tends to increase to 24.75 ± 5.5 seconds in larvae from the same reproduction conditions but developed at elevated temperature (n = 10), although not significantly (Student’s t-test, p-value = 0.089). This trend of longer lasting brain activity response was also observed in offspring of parents reproduced at elevated temperature but reared in control conditions (duration = 23.65 ± 5.6 seconds, n = 10) even though it was again not statistically significant (Student’s t-test, p-value = 0.234). However, the brain response to CAC of offspring from parents reproducing and reared at elevated temperature lasted 18.48 ± 7.2 seconds (n = 9), which is more similar to control.

We verified that the telencephalon area (11 010.65 ± 1111.89 µm^2^; Table S2) did not vary due to parental spawning temperature (two-way ANOVA, p-value = 0.782), nor offspring temperature (two-way ANOVA, p-value = 0.648) nor the interaction between those two factors (two-way ANOVA, p-value = 0.257; Fig. S4). This suggests that the differences observed are not due to physiological development accelerated by elevated temperature in the forebrain.

## Discussion

We explored how near-future warming experienced within and between generations affects the neuronal response of zebrafish larvae as they sense an alarming olfactory stimulus. We found that parental spawning and early embryogenesis in elevated temperature accelerated the time of maximal activity response in the telencephalon of the offspring, regardless of their own rearing temperature. Furthermore, brain activity duration was affected by the interaction between parental spawning and offspring developmental thermal conditions.

Larvae from parents reproducing at elevated temperature reached their maximum brain activity response to CAC earlier than their half-siblings arising from reproduction at control temperature. Although there is previous evidence that offspring can inherit neural activity traits from their parents as they sense olfactory stimuli (Liu *et al*., 2017), our results show that during this response neuron excitability is also increased by the thermal environment of their parents at the time of reproduction. Furthermore, it occurs through different mechanisms than the known direct effect of water warming on axon conductance (Flerova and Gdovskii, 1975; Døving and Belghaug, 1977) as we observed a faster brain activity maximum regardless of the offspring’s developmental temperature. There are different pathways through which brain activity response could be changed after parental spawning in elevated temperature. First, such modifications could be non-genetic inheritance from either the paternal, maternal or both origins involving molecular regulation of neurodevelopment (Colson *et al*., 2019; Chan *et al*., 2020). Another potential mechanism may be that elevated temperature at reproduction directly acts on gametes or even non-sperm components of ejaculate, as seen in another fish species where sperm thermal environment influenced post-hatching performance (Läinen *et al*., 2018). In the latter case, cellular stress responses to warming in various components of the semen (Lane *et al*., 2014; Menezo *et al*., 2016) were previously documented and could also be responsible for epigenetic modifications (Immler, 2018) causing the observed changes in the offspring. Finally, a faster time of maximum brain activity could be a phenotypic plastic response expressed by the offspring itself triggered at their very early embryonic stage, due to their exposure to parental thermal conditions then. In other fishes, an increase in incubation temperature during gastrulation provoked changes in hatchling phenotype, revealing a critical window of development (Mueller *et al*., 2015; Melendez and Mueller, 2021) which shows that thermal experience during early embryogenesis provides relevant information about later conditions in fish’s life (Fawcett and Frankenhuis, 2015). Here, it could be that the parental environmental conditions influenced the offspring directly right after fertilization, during the short time before fertilized eggs were exposed to their developmental temperature during their week of incubation. Therefore, whether through parental effects, or abiotic effects on the gametes, sperm or fertilized eggs, modifications leading to changes in neuron activity are caused by the parental thermal environment.

A possible target of temperature-driven modifications in the zebrafish offspring may be processes involved in excitatory synaptic transmission. Particularly, the dopaminergic system may be involved in intergenerational effects of warming on zebrafish brain activity: dopaminergic neuron populations are found in the larval zebrafish telencephalon, notably in but not restricted to the olfactory bulb (McLean and Fetcho, 2004). Furthermore, dopaminergic neurotransmission influences odour responses (Bundschuh *et al*., 2012) and dopamine modulates neuron firing (Schärer *et al*., 2012; McGregor and Nelson, 2020). Finally, the dopaminergic system can be affected in a transgenerational way through modified expression of dopaminergic receptors (Yu *et al*., 2021), which is also sensitive to warming in fish developing stages (Giroux, Gan and Schlenk, 2019). Here, epigenetic modification of dopaminergic receptors expression in the telencephalon caused by parental temperature conditions could increase neuron excitability in the offspring, causing repeated firing during CAC detection reaching a faster time of maximal activity. Another possibility is that parental exposure to elevated temperature could change molecular processes involved in inhibitory transmission, such as GABAergic neurotransmission. GABA is expressed as an inhibitory neurotransmitter in the majority of interneurons in the olfactory bulb even at the larval stage (Miyasaka *et al*., 2013). In fish, proteins and genes involved in the transport and functioning of GABA were found altered within and across generations after exposure to another environmental stressor, ocean acidification, with consequences on signalling and antipredator behaviour (Schunter *et al*., 2016; Williams *et al*., 2019). Furthermore, elevated temperature has the potential to alter GABAergic-mediated behaviours (Clements, Bishop and Hunt, 2017). Here, warming experienced by the parents could affect GABAergic interneurons in a way that would reduce their inhibitory input on the olfactory circuitry, making neuronal populations more excitable, and causing them to reach a maximum activity level sooner than in control conditions. Overall, our results support a model in which neural excitability in the telencephalon of zebrafish larvae is affected by elevated temperature, with several possible and non-exclusive mechanisms of non-genetic inheritance at play either within or across generations.

Although not significant, development at higher temperature seemed to increase response duration to CAC in the telencephalon, consistent with previous research finding positive correlations between neural activity and temperature (Beltrán *et al*., 2021). Larvae whose parents reproduced at elevated temperature yet developed in control conditions tended to have a longer lasting response, suggesting that elevated temperature during early life but also at the time of breeding can prolong the brain activity response of larvae to an alarm cue. Longer responses may occur through increased influx of calcium which would enhance neurotransmitter release in synaptic areas after exposure to higher temperature (De Boeck *et al*., 1996; Peng *et al*., 2007; Vargas-Chacoff *et al*., 2019). Alternatively, high intracellular calcium levels may be maintained longer because other calcium signals are involved, regulating additional processes such as gene transcription (Berridge, Bootman and Roderick, 2003; Brini *et al*., 2014). In both cases, synaptic transmission modifications would likely result from longer activity of the olfactory circuitry, a phenomenon known as synaptic plasticity. Therefore, our results are consistent with early developmental temperature influencing synaptic plasticity processes in fish (Dunlap, 2016; Bernal *et al*., 2022), as seen in other ectothermic species or stages (Groh, Tautz and Rössler, 2004; Peng *et al*., 2007; Bertin *et al*., 2018) and those neural circuits could produce modifications in behaviours with warming (Tautz *et al*., 2003; Bertin *et al*., 2018; Beltrán *et al*., 2021).

Interestingly, there are complex effects of temperature on brain signal duration depending on the time of exposure. While exposure to warming during spawning and early embryogenesis produced a longer lasting response in control offspring, when the offspring developed in elevated temperature a “restoration” of brain activity duration similarly to control was observed. This may represent an anticipatory parental effect, which consists in adjusting the offspring’s phenotype to the parental thermal environment, under the assumption that the parental environment corresponds to that of the offspring (Burgess and Marshall, 2014; Bale, 2015). Since “restoration” of brain activity duration was only observed when both parents and offspring were kept at the same elevated temperature, exposure to parental conditions may be effective to rescue normal brain functions when they would otherwise be affected as seen in another ectotherm species (Arai *et al*., 2009) but provided that the parental thermal environment is a reliable predictor of the offspring’s rearing temperature (Donelson *et al*., 2018). For stenothermic aquatic species the temperature experienced during parental reproduction may correspond well to the temperature experienced by the next generation (Somero, 2010; Logan and Buckley, 2015), however zebrafish are subject to a wide range of seasonal thermal variation in their natural environment (Spence *et al*., 2008; López-Olmeda and Sánchez-Vázquez, 2011). Overall, elevated temperature would tend to increase the brain activity response duration during olfactory stimulation which could have important potential implications on olfactory-mediated behaviour. However, this effect can be compensated through parental anticipatory effects, suggesting that the impact of near-future predicted thermal conditions may be limited owing to exposure to the parents’ environment.

Nevertheless, the adaptive value of such parental conditioning effects conferred to the offspring is not clear, and it is often difficult to assess whether the response to parental environments is adaptive (Donelson *et al*., 2018; Candolin and Rahman, 2023), where changing one’s phenotype can buffer against effects of environmental change. Although elevated temperature experienced as embryos was shown to have persistent positive effects on performance at elevated temperature for zebrafish adults (Scott and Johnston, 2012), such developmental temperature acclimation cannot always be expected as beneficial (Leroi, Bennett and Lenski, 1994; Huey *et al*., 1999). When it comes to plasticity observed at the brain level, warming in goldfish also caused hyperexcitability and these neural changes resulted in a faster escape response, but decreased the directionality of the escape (Szabo *et al*., 2008), suggesting that warming could have mixed effects on antipredator behaviour. Therefore, it is unclear whether parental conditioning effects on the time of maximum brain activity observed in our study would prove advantageous to the offspring. Nevertheless, previous research on other vertebrates showed that timing of olfactory processing is important for odorant recognition, as perception depended on activation latencies of brain regions and cells, especially the cells that are activated earliest (Chong *et al*., 2020). This suggests that faster activation of the telencephalon caused by the parental environmental conditions could influence the perception of CAC and potentially the antipredator behaviour triggered by it (Speedie and Gerlai, 2008). Furthermore, one may view as adaptive the restoration of brain activity duration when both generations experience elevated temperature, given that it maintains the phenotype observed in control conditions.

In summary, our findings show an effect of future-predicted thermal conditions on developing zebrafish, reaching maximal telencephalic activity sooner when exposed to an olfactory alarm cue. Furthermore, the thermally induced prolongation of neural activity could be mitigated if parents reproduced at such elevated temperature, suggesting potential parental anticipatory effects or early developmental plasticity. These effects, particularly during sensitive early life-history stages, may serve to enhance persistence and adaptability in the face of environmental change provided that parental conditions accurately predict the offspring’s environment.

## Author contributions

JMS conceived and carried out the experiments with input from CS. JLS provided the breeding animals and input on zebrafish breeding. JMS analysed the data under the supervision of CS. JMS wrote the paper with input from CS and JLS.

## Acknowledgments

JMS and this study were funded by the start-up of CS from the University of Hong Kong. The study was partially funded by a General Research Fund (17300721) and the NSFC Excellent Young Scientist Award (AR225205) to CS. We thank Yan Chit Kam who contributed to the daily care of breeding animals and Helen Leung who helped with confocal imaging and all the members of the lab for support.

